# Interaction between gene expression and morphokinetic parameters in undisturbed human embryo culture

**DOI:** 10.1101/2024.06.04.596054

**Authors:** Hui Xiao, Adam Stevens, Helen L. Smith, Karolina Szczesna, Maria Keramari, Gregory Horne, Andras Dinnyes, Susan J. Kimber, Pietro Lio, Daniel R. Brison

## Abstract

The clinical in vitro fertilisation (IVF) need for optimal culture conditions and improved selection of human embryos for transfer led to the development of time-lapse systems built into incubators to allow a stable, well-defined and undisturbed culture environment with continual monitoring of embryo development. Clinical data suggest that both undisturbed culture of embryos and selection algorithms based on time-lapse morphometric parameters can potentially improve embryo development and live birth rates. However, there remains a need to validate and understand the mechanisms underlying the potential benefits of the time-lapse technology in IVF. In this study, we compared the gene expression of human embryos grown in an undisturbed time-lapse system with a conventional incubator and showed that there is no fundamental difference in the developmental program of the undisturbed embryos, which provides important reassurance concerning the time-lapse technology. We then performed a network-based integrative analysis based on the undisturbed blastocyst transcriptomes and identified time-lapse parameter correlated genes. These showed cross talk with identified embryo development gene functional modules, suggesting potential molecular mechanisms underlying the clinical predictive capabilities of embryo time-lapse morphokinetic parameters for subsequent live birth. This study suggests methodologies for assessing the impact of additional predictive correlates of value for optimal embryo development and understanding their mechanisms of action.

## Introduction

In vitro fertilization (IVF) is a well-established method of infertility treatment and more than 8 million IVF-conceived children have been born worldwide since the first in 1978 (1). However, success rates still remain relatively low, the IVF laboratory environment is not fully optimized, and objective information on human embryo quality and methods for embryo selection are still lacking. Time-lapse (TL) technology is a relatively new method for embryo selection and has proven amenable to integration into IVF laboratories around the world. The clinical need for optimal culture conditions and continual monitoring of human embryo development led to the development of TL systems built into incubators that ensures a stable, undisturbed culture environment without the removal of embryos from the incubator for observation and without group culture of more than one embryos in culture drops and the interference of paracrine embryo secreted factors. The use of TL incubators can improve embryo survival, clinical pregnancy and live birth rates compared to standard incubators (2–4). In a cohort study performed in our centre, TL embryo culture was associated with an increased live birth rate (odds ratio of 1.43) compared to a high-quality benchtop incubator (5).

In addition to undisturbed and individual embryo culture, TL monitoring offers a more objective assessment of embryos for an improved selection process, compared to conventional embryo morphology, which encompasses only intermittent snapshots of a complex and dynamic process. TL analysis provides a multitude of informative parameters at each stage of embryo development including pronuclear (PN) formation, cleavage, symmetry, timing of cell divisions and timing and quality of blastocyst formation. While the sheer volume of data provided has proven challenging to develop into embryo selection algorithms which demonstrate clear clinical benefit (6), a number of different scores based on the various kinetic parameters have been described and may have some predictive value for embryo selection.

Thus, although TL has already been integrated into many laboratories around the world, an outstanding need to validate and understand the benefits remains, as for all new technologies entering ART (7). We introduced TL technology into Saint Mary’s Hospital IVF clinic in a two-step process; the pre-clinical scientific study described here, followed by a clinical validation study now published as Kalleas et al (5).

Therefore, the aims of this study were, firstly, to compare blastocysts developed in a TL incubator to those grown in a standard “box” incubator, using embryo whole transcriptome gene expression profiling to provide a readout of any differences in embryo development. Secondly, we proposed a network-based integrative analysis framework to investigate the associations between TL parameters and the blastocyst transcriptome, to explore at a molecular level the mechanistic basis of the predictive capabilities of TL parameters for optimal embryo development.

## Materials and Methods

### Human embryo culture

All human embryos were donated to research with approval from Central Manchester Research Ethics Committee and the Human Fertilisation and Embryology Authority (research license R0026). Embryos from Saint Mary’s Hospital and Manchester Fertility, Manchester, were frozen at the 2PN stage and thawed using the Embryo Freezing Pack (Medicult UK Ltd., Surrey, UK). Thawed embryos were placed into incubators and cultured to the blastocyst stage in G1 medium (Vitrolife, Goteborg, Sweden) until day 3, followed by G2 medium (Vitrolife) from day 3 to blastocyst stage on day 6 at 37°C, 6% CO_2_. Four of the blastocysts developed in a high quality conventional embryo culture CO2 incubator (Heracell; Hereaus, UK), in humidified atmospheric oxygen, with daily removal of embryos for morphological observation as per standard practice. These are referred to hereafter as Standard Incubation (SI) blastocysts; this reference transcriptome dataset has been deposited in the GEO repository (GSE110693) (8). The other 10 blastocysts developed in the EmbryoScope™ incubator (Fertilitech) at 5% O2, with remote imaging allowing embryos to remain in undisturbed culture. These 10 blastocysts were selected for transcriptome analysis to match the 4 SI blastocysts in the reference dataset, by day of blastocyst formation (day 6) and morphological grade, and are hereafter referred to as EmbryoScope (ES) blastocysts. The EmbryoScope records images every 5 minutes, providing video data for TL analysis for the 10 ES blastocysts. The transcriptome data of the 10 ES blastocysts have been deposited in the GEO repository (accession GSE180605).

### Time-lapse parameters

Time-lapse technology with image analysis software allows for the tracking of specific timings between developmental events, known as TL morphokinetic parameters. TL data are used to identify pronucleus fading (PNF), also known as syngamy, as well as the timing of the first cell division to two cells (T2), followed by further divisions corresponding to three (T3), four (T4) and five cells (T5). Four TL parameters, T5, S2, CC2 and CC3, have been identified as the most important for human embryo development (2, 9, 10). T5 is an annotation point in which the embryo finishes the division to become five cells. S2 is the synchrony of the division from two cells to four cells (T4-T3). CC2 is the time of second cell cycle, which corresponds to the duration of the 2-blastomere embryo stage (T3-T2) period. CC3 is related to the third cell cycle or, more precisely, the time interval between the 3-cell and 5-cell stages (T5-T3). Morphokinetic characteristics have been reported to be correlated with implantation possibility (4) and many morphokinetic classification models have been developed based on TL parameters for embryo selection. Cruz et al. (11) proposed a hierarchical classification method based on T5 and S2 to classify embryos into four morphokinetic categories, A, B, C and D, which predict the embryos’ implantation potential decreasingly (Figure S1).

### Blastocyst grading

Blastocysts were graded depending on three parameters: the expansion status (EXP), the inner cell mass (ICM) and the trophectoderm (TE) (12). Each of these features is given a numerical score on a scale of 1-3, with increasing score reflecting increasing quality, based on: degree of expansion (EXP), the size, number of cells, organisation and integrity of the Inner Cell Mass (ICM), and the number of cells and degree of organisation of the Trophectoderm Epithelium (TE).

### Blastocyst gene expression

#### Embryo preparation for global amplification (PolyAPCR) - Affymetrix® Gene Profiling Array cGMP U133 P2

PolyAPCR (13) was used to amplify all polyadenylated RNA in a single embryo. Embryos were lysed and then reverse transcribed as previously described (8, 13, 14). PCR amplification of the polyAcDNA product was performed by two linked sets of 25 cycles. The first 25 cycles were denatured at 94 °C for 1 minute, followed by primer annealing for 2 minutes at 42 °C and 6 minutes of extension at 72 °C. The second 25 cycles were denatured at 94 °C for 1 minute, followed by primer annealing at 42 °C for 2 minutes and extension at 72 °C for 1 minute. A second round of amplification using EpiAmp™ and biotin-16-dUTP labelling using EpiLabel™ was performed in the Paterson Cancer Research Institute Microarray Facility, as previously described (8, 15). For each sample, our minimum inclusion criterion was the expression of β-actin evaluated by gene specific PCR. Labelled PolyAcDNA was then hybridised to the GeneChip® Human Genome U133 Plus 2.0 Array (Affymetrix, SantaClara, CA, USA) that covers over 47,000 transcripts and have been commonly used for human transcriptome analysis, and data initially visualised using MIAMIVICE software, as described before (15).

#### Pre-processing of microarray data

The statistical and graphical R computing language (16) was used together with Bioconductor packages (17), to assess quality control of microarray data (Array Quality Metrics package) (18) and to pre-process the raw data. The raw microarray CEL files were processed and normalised by using the Robust MultiArray (RMA) normalization (19). Detection calls were performed on the normalised data by using the mas5calls (20) function provided by the R package affy (21), which indicate whether a transcript is detected in a sample as present (P), absent (A), or marginally-present (M).

#### Statistical analysis

Unsupervised hierarchical clustering and principle component analysis (PCA) were applied on the gene expression data of the SI and ES blastocysts. Empirical Bayes moderated t-test proposed by Limma package (22) was used to assess differential expression of genes between SI and ES blastocysts. P-values of differential expression were adjusted by Benjamini-Hochberg False Discovery Rate (FDR) correction (23). Genes with FDR lower than 0.05 were defined as DEGs.

#### Gene Ontology enrichment

To evaluate the functional involvement of the TL parameter correlated genes, we performed Gene Ontology (GO) (24) enrichment based on the hypergeometric test. The significant GO biological processes with FDR lower than 0.05 were selected as the enriched function terms for the DEGs.

### Network-based integrative analysis framework for studying associations between TL parameters and preimplantation development

We proposed a network-based integrative analysis framework, illustrated in Figure 1, to study the associations between TL parameters and human preimplantation embryo development by selecting TL parameter correlated genes and their associations with preimplantation embryo development.

**Figure 1.**
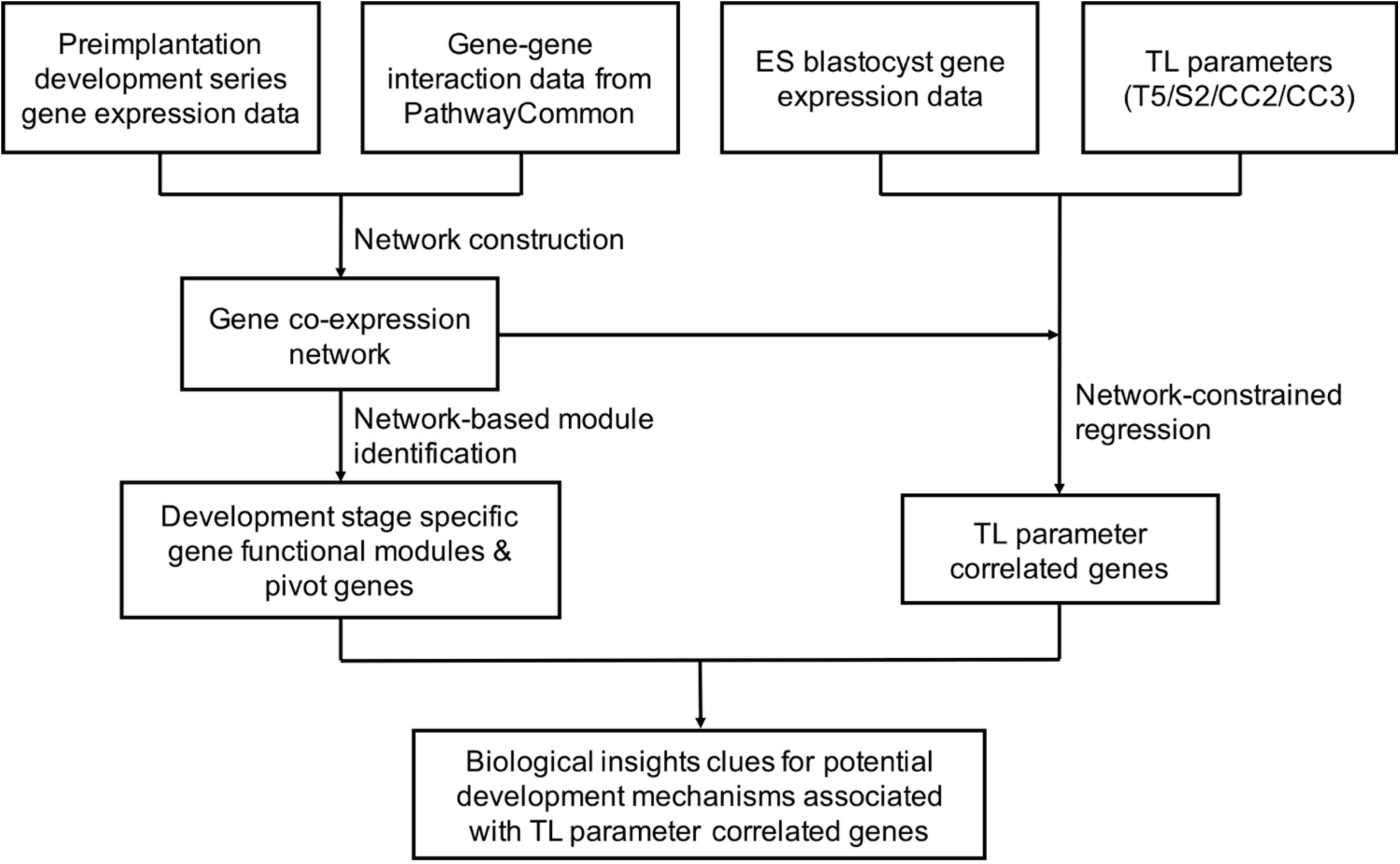
Flowchart of the network-based integrative analysis framework.

#### Construction of gene co-expression network of human early embryo development

We constructed a gene co-expression network across human preimplantation embryo development by integrating a gene-gene functional interaction (GGI) network derived from the public database PathwayCommon (version 8) (25) with the previous in-house generated microarray profiling gene expression data of human embryos across multiple developmental stages, including oocyte, 4-cell, 8-cell and blastocyst stage (8). For each linked gene pair in the GGI network, the co-expression between the two genes was assessed by Pearson Correlation Coefficient (PCC). The PCC values were assigned as weights to the edges of GGI network, and we therefore obtained the weighted co-expression gene network of embryo development (hereafter referred to as “co-expression network”).

#### Selection of TL parameter correlated genes from blastocyst gene expression using network-constrained regression method

TL parameter correlated genes are defined as those whose expression in blastocysts is correlated with a specific TL parameter (T5, S2, CC2 or CC3). TL parameter correlated genes are selected by using network-constrained regression. We proposed a network-constrained regression method based on an existing network-constrained regression method that was implemented by using Generalized Boosted Lasso (GBL) algorithm (26). We modified the GBL network-constrained regression model by incorporating the edge weights of the network into regularization when estimating the regression coefficients, and thus the proposed method is called edge-based GBL (eGBL) network-constrained regression. We applied eGBL on blastocyst gene expression data to select TL parameter correlated genes for each TL parameter.

Suppose we have *n* embryos. Let *y* = (*y*_1_, …, *y_n_*)*^T^* be a vector containing the values of a TL parameter of *n* blastocysts. Let *x_i_* = (*x_i1_*, …, *x_ip_*) be a vector containing the expression values of *p* genes in the blastocyst *i*, corresponding to the TL parameter value *y_i_*, where *i* = 1, …, *n*. The simple linear model for the TL parameter and blastocyst gene expression is defined as:

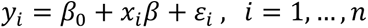

Where *β* = (*β*_1_, …, *β_p_*)*T*represents the coefficients corresponding to the *p* genes and *β*_0_ is the intercept of the linear model. *ε_i_* is an unobserved random variable of the systematic error. A regression coefficient *β_j_* (*j* = 1, …, *p*) can be interpreted as the degree of association between the expression of gene *j* and the TL parameter *y*. A positive *β_j_* indicates the correlation between gene *j* and the TL parameter, while a negative *β_j_* indicates the anti-correlated relationship.

To avoid the overfitting in the linear models for coefficient estimation, *β* was estimated by regularized regression model fitting procedure which performs Ordinary Least Square (OLS) estimation by adding a penalty function to the loss function to constrain the values of coefficients. The loss function is defined as

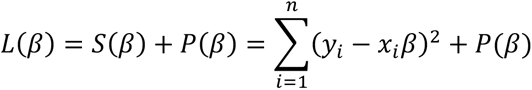

The estimated coefficients are given by minimizing *L*(*β*).

In order to incorporate the prior knowledge of the correlation relationships between genes, we defined the penalty function based on a prior gene co-expression network which provides dynamic functional correlations between genes during human preimplantation embryo development. The penalty function is given by

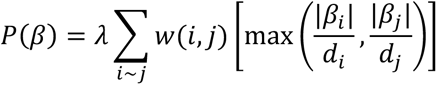

where *W*(*i*, *j*) represents the co-expression weight of linked gene *i* and gene *j* in the network. *d_i_* and *d_j_* represent the weight summary of all linked nodes in the network for gene *i* and gene *j* respectively.

#### Identification of key genes and gene functional modules from preimplantation development co-expression network

In order to explore the involvement of TL parameter correlated genes in human preimplantation embryonic development, we performed a system biology analysis based on the co-expression network to identify key genes and gene functional modules associated with specific development stages. The co-expression network was filtered by only keeping the significant co-expression edges with p-values of PCC lower than 0.05. Co-expressed gene modules were detected from the network by using OCG algorithm (27) and modules containing a minimum of 5 genes were retained. To identify associations between gene modules and development stages, we first filtered gene modules that were significantly enriched with high variance of gene expression across all development stages by gene set enrichment analysis (p-value < 0.05) (28), and then identified gene modules associated with specific development stages by using group Lasso binomial regression (29). Genes located between modules in the network were defined as inter-modular genes. If an inter-modular gene linked to more than one module with at least two connections to each module, it is referred to as a pivot gene.

## Results

### Gene expression of human embryos developed in EmbryoScope (ES) and Standard Incubation (SI) systems

To investigate whether time-lapse incubation is associated with altered gene expression patterns of human embryos, we performed statistical analysis on the in-house generated transcriptome data of 10 ES developed blastocysts and our previously published transcriptome data of 4 SI developed blastocysts (8), including unsupervised hierarchical clustering, principle component analysis (PCA) and differential gene expression analysis. The raw microarray data of the 10 ES blastocysts and the 4 SI blastocysts were pre-processed by using RMA and the detection calls were obtained using mas5call. A gene was defined as expressed in one of the two groups if it was detected as “present” in at least 75% (3/4 or 8/10) of the embryos. By this definition, 2920 genes were expressed in either the SI or ES blastocyst groups. Unsupervised hierarchical clustering was performed on the gene expression data of the two groups of blastocysts. The gene expression pattern was not different between the ES and SI groups and the embryos were classified into mixed clusters, with SI blastocysts tending to be clustered more closely with ES blastocysts than with other SI blastocysts (Figure 2A). Principle component analysis (PCA) also did not identify two distinct clusters for ES and SI groups along the first two principal components (Figure 2B). Differential gene expression between the ES and SI blastocysts was evaluated by using Limma package and only 40 out of the 2920 genes (1.4%) were identified as differentially expressed genes (DEGs) at FDR 5% between the ES and SI groups (Figure S2).

**Figure 2.**
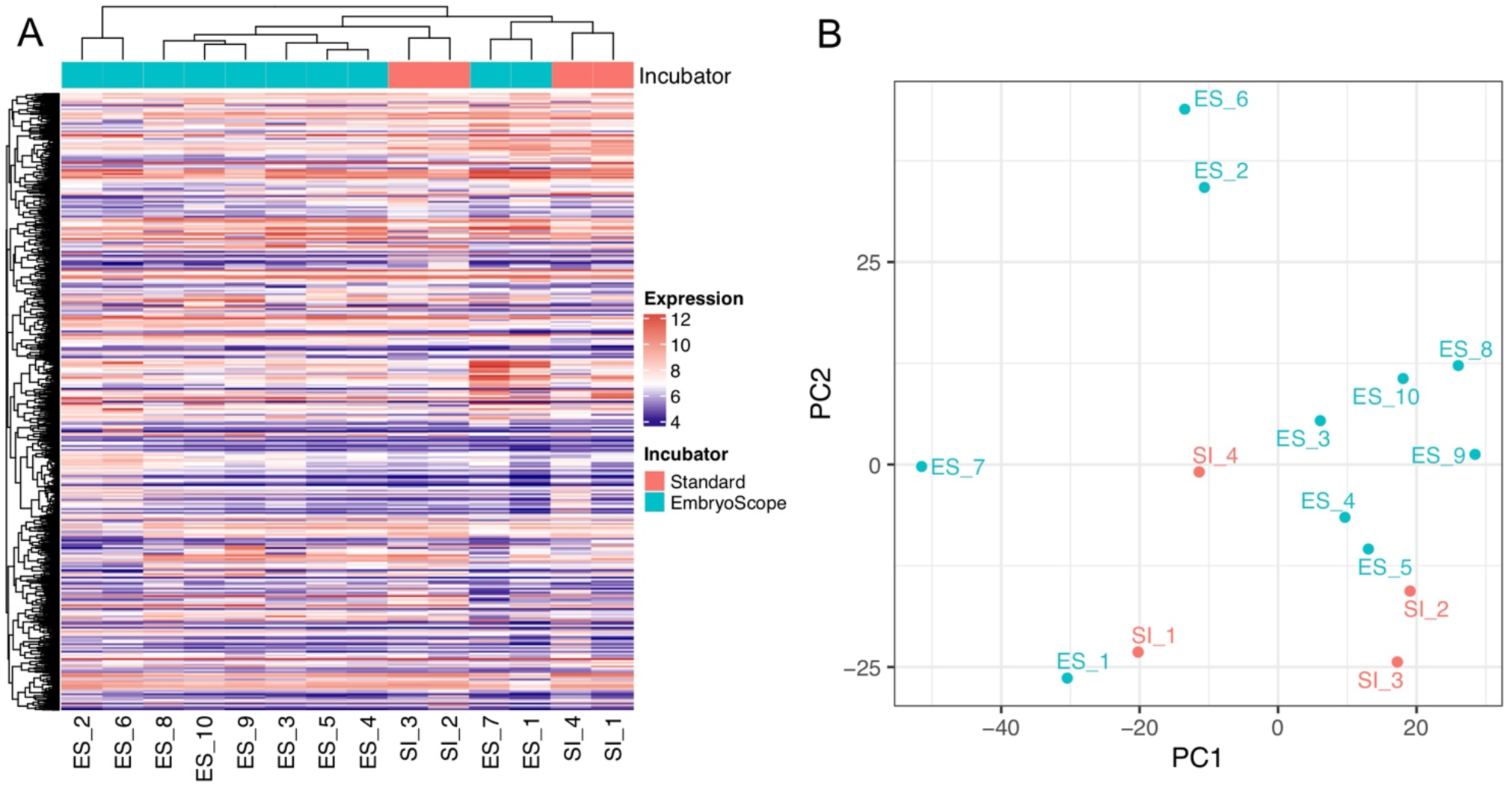
Hierarchical clustering (A) and PCA (B) of embryos developed in the standard incubation (SI) and the EmbryoScope (ES) systems based on the blastocysts gene expression.

### Identification of TL parameter correlated genes

In this study, TL parameters were obtained for the 10 ES blastocysts, which provided the opportunity to explore the potential molecular mechanisms associated with the predictive capabilities of TL parameters for embryo development and live birth, by integrating the blastocysts transcriptome data. To investigate the associations between TL parameters and the blastocyst transcriptome, we proposed a network constrained regression method eGBL for selecting TL parameter correlated genes, which takes into account the prior defined network structured correlation relationship between genes. Comparing the performances of eGBL with several common regression methods on three simulated datasets, we found that eGBL offers better results in terms of feature selection than other common methods such as Lasso (30), Elastic Net (31), Grace (32) and GBL (26) (Figure S3).

TL parameter correlated genes were selected using eGBL by incorporating a gene co-expression network of human preimplantation early embryonic development. The co-expression network was constructed by integrating the gene-gene interaction network derived from PathwayCommon with gene expression data of human embryos across multiple preimplantation development stages (8). The raw gene expression data were pre-processed following the same procedures for the 10 ES blastocysts data. The co-expression correlation coefficient evaluated by Pearson Correlation Coefficient was assigned to each interacted gene pair (i.e. each edge) in the gene-gene interaction network. The co-expression network consists of 12,961 genes and 493,759 interactions.

Because the number of genes is exponentially larger than the number of blastocysts, we firstly screened the genes to reduce the dimension. For each TL parameter, the correlation between the parameter and each gene’s expression was evaluated by the Spearman’s Rank Correlation Coefficient (SCC). The genes significantly correlated with the parameter (SCC p-value < 0.05) were selected as the candidate genes. TL parameter correlated genes were selected for each of the four TL parameters, T5, S2, CC2 and CC3, by applying eGBL on gene expression data of the candidate genes. Totally 68 genes were identified correlated with the TL parameters (Figure 3 and Table 1). Most genes were correlated with a single TL parameter and very few overlapped between different TL parameters (Figure S4). Several TL parameter correlated genes have been reported to play important roles in mammalian embryonic development, e.g. T5 correlated genes CALCOCO2 (33), HSF2 (34), MAGE11 (35) and EGFR (36), S2 correlated genes HAND1 (37) and GRHL2 (38), CC2 correlated genes CDX2 (39) and PARP1 (40), CC3 correlated gene PPARD (41).

**Figure 3.**
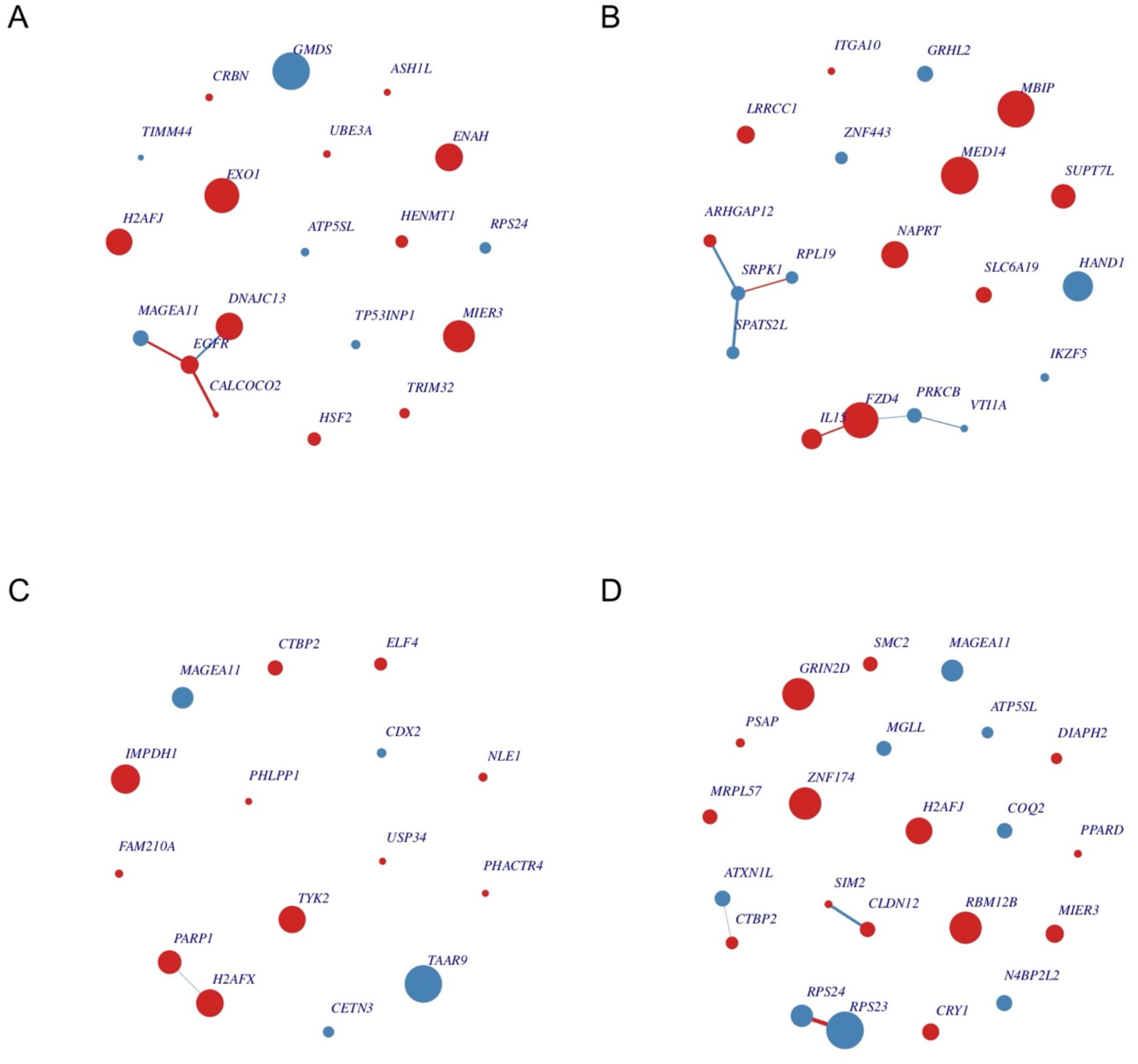
TL parameter correlated genes: (A) T5, (B) S2, (C) CC2 and (D) CC3. Each node represents a gene and the node size is proportional to its weight of regression coefficient. Nodes in red represent genes whose expression is positively correlated with the TL parameter, while nodes in blue represent genes negatively correlated with the TL parameter. Each edge represents the co-expression interaction between two genes and the edge width is proportional to the correlation coefficients. Edges in red represent the positive co-expression and edges in blue represent the negative co-expression.

**Table 1.** TL parameter correlated genes and enriched biological processes.

| TL parameter | TL parameter correlated genes | Enriched biological processes (top 5 most significant) |
| --- | --- | --- |
| T5 | GMDS, EXO1, MIER3, ENAH, DNAJC13, H2AFJ, EGFR, MAGEA11, HSF2, HENMT1, RPS24, TRIM32, TP53INP1, ATP5SL, CRBN, UBE3A, ASH1L, CALCOCO2, TIMM44 | • response to UV<br>• regulation of catabolic process<br>• RNA metabolic process<br>• nucleobase-containing compound metabolic process<br>• positive regulation of transcription, DNA-templated |
| S2 | MED14, MBIP, FZD4, HAND1, NAPRT, SUPT7L, IL15, LRRCC1, SLC6A19, GRHL2, PRKCB, SRPK1, SPATS2L, ARHGAP12, ZNF443, RPL19, IKZF5, VTI1A, ITGA10 | • developmental process involved in reproduction<br>• nucleobase-containing compound biosynthetic process<br>• positive regulation of macromolecule biosynthetic process<br>• heterocycle biosynthetic process<br>• regulation of sequence-specific DNA binding transcription factor activity |
| CC2 | TAAR9, IMPDH1, H2AFX, TYK2, PARP1, MAGEA11, CTBP2, ELF4, CETN3, CDX2, NLE1, FAM210A, PHLPP1, PHACTR4, USP34 | • regulation of signal transduction<br>• single-organism developmental process<br>• developmental process<br>• regulation of response to stimulus<br>• regulation of signaling |
| CC3 | RPS23, ZNF174, GRIN2D, RBM12B, H2AFJ, RPS24, MAGEA11, MIER3, CRY1, N4BP2L2, ATXN1L, COQ2, CLDN12, MGLL, MRPL57, SMC2, CTBP2, ATP5SL, DIAPH2, PSAP, PPARD, SIM2 | • negative regulation of transcription from RNA polymerase II promoter<br>• negative regulation of transcription, DNA-templated<br>• negative regulation of nucleic acid-templated transcription<br>• negative regulation of RNA biosynthetic process<br>• negative regulation of RNA metabolic process |

Gene Ontology biological processes significantly enriched with TL parameter correlated genes suggested potential associations between TL parameters and a number of crucial functions were related to embryonic development such as regulation of transcription, cell cycle, metabolic process, viral life cycle, signal transduction and histone modification (Table 1 and Figure S5-S8).

### Association between TL parameter correlated genes and blastocyst qualities

The 10 ES blastocysts were allocated into 4 morphokinetic categories based on T5 and S2 following the classification method proposed by Cruz et al. (11) (Figure S1). Unsupervised clustering was performed on the blastocysts based on expression of the identified TL parameter correlated genes (Figure 4) to explore the correlations between TL parameter correlated gene expression and morphokinetic categories. Blastocysts with higher qualities (i.e. higher blastocyst grades) tended to be grouped together by TL parameter correlated gene expression. The blastocysts ES_1, ES_4, ES_5, and ES_7 were clustered into a group because of highly correlated expression in TL parameter correlated genes. The group exhibited higher blastocyst qualities (average grades 3.8, 2 and 2.3 for EXP, ICM and TE) than the other embryos (average grades 2.2, 1.3 and 0.8 for EXP, ICM and TE). The proportion of morphokinetic category A (i.e. predicting the highest implantation potential) in this group (50%) was higher than in the other embryos (16.7%). The concordance of the gene expression, blastocysts qualities and morphokinetic categories suggests the TL parameter correlated gene expression correlated with implantation potential.

**Figure 4.**
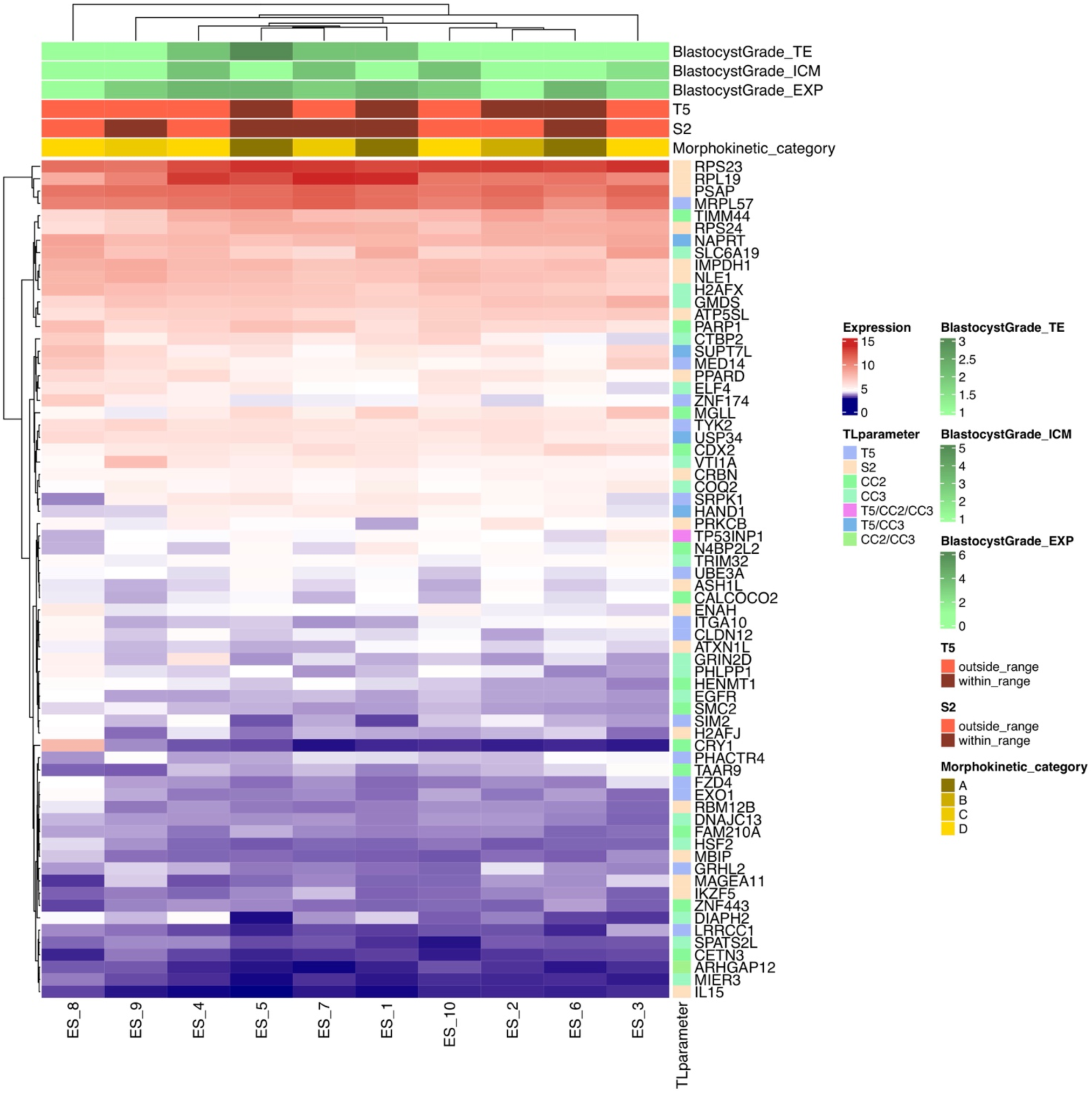
Hierarchical clustering of blastocysts based on TL parameters correlated gene expression. The heatmap presents the bi-clustering of both genes (by rows) and ES blastocysts (by columns). The colored bars represent blastocyst categories according to blastocyst grades, TL parameter optimal ranges, morphokinetic categories proposed by Cruz et al. (11) (Figure S1), and the associated TL parameters.

Based on the TL parameter correlated gene expression, we classified the 10 ES blastocysts into two groups by using unsupervised hierarchical clustering: the good-quality group consisting of ES_1, ES_4, ES_5, and ES_7, which exhibited higher blastocyst grades, and the poor-quality group consisting of the other 6 blastocysts that exhibited lower blastocyst grades. Several TL parameter correlated genes have been reported to be related with peri-implantation development, including CDX2, GRHL2, HAND1, PARP1 and PPARD, which play important roles in trophectoderm development, trophoblast differentiation and placentation. Expression of HAND1 was significantly lower in poor-quality blastocysts compared to the good-quality blastocysts (Figure 5). There were also marked differences in gene expression of PARP1 and PPARD between the good-quality and poor-quality groups, although no significance was observed due to the small sample size (Figure 5). The gene expression of the peri-implantation markers suggests potential correlation between the TL parameters S2, CC2 and CC3 and morphokinetic categories as well as the implantation potential.

**Figure 5.**
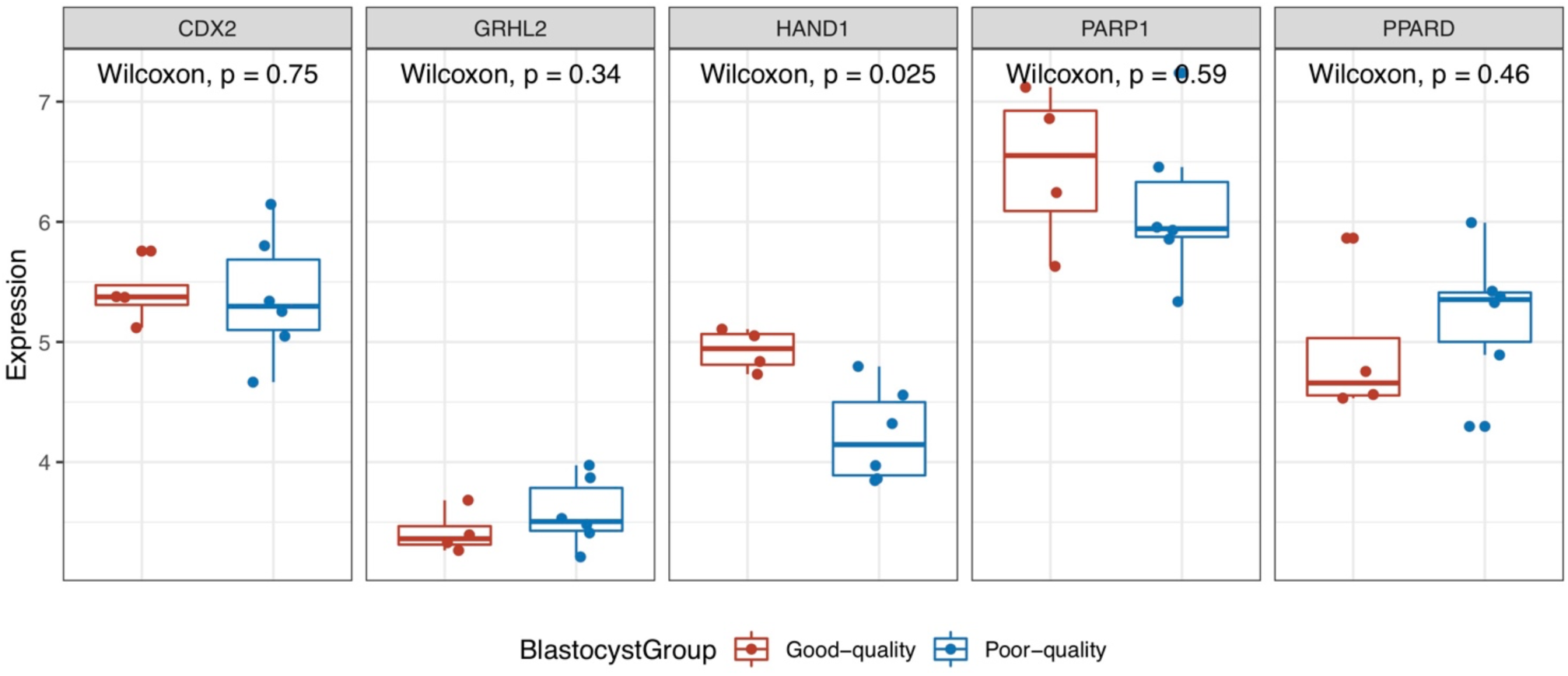
Expression of 5 peri-implantation development related genes between good-quality and poor-quality ES blastocyst groups.

### TL parameter correlated genes in human preimplantation embryonic development

From the co-expression network, 42 functional gene modules were identified as embryonic development stage-specific modules, with 14, 17, 16 and 14 modules being specific for oocyte, 4-cell, 8-cell and blastocyst stages, respectively (Table S1). The largest module consists of 42 genes, while the smallest one includes 5 genes. Through functional enrichment analysis with GO, we found that all 42 modules were significantly enriched with GO biological processes.

To explore the involvement of TL parameter correlated genes in human preimplantation embryonic development, we annotated them to the co-expression network of human embryonic development, with reference to the 42 development stage specific modules. In the co-expression network, genes located between the 42 stage-specific modules are referred to as inter-modular genes. If an inter-modular gene has at least two links to genes belonging to a module, the inter-modular gene is defined as crosstalking with the module. Such crosstalk between genes and modules suggest potential functional associations between them. The inter-modular genes crosstalking with more than one module were selected as pivot genes (Table S2). Based on the stage-specific modules and the pivot genes, we annotated the TL parameter correlated genes in the co-expression network and identified several key genes with important topological characteristics (Table 2). The key topological characteristics of the TL parameter correlated genes in the co-expression network suggest potential critical roles in human preimplantation embryonic development. They also provide clues for exploring the underlying molecular mechanisms that account for the association between early stage TL parameters and blastocyst development.

**Table 2.** TL parameter correlated genes with key topological characteristics in human preimplantation embryonic development gene co-expression network.

| Gene | TL parameter | Topological characteristics | Embryo stages |
| --- | --- | --- | --- |
| EGFR | T5 | Intra-modular gene | 4-cell |
| EXO1 | T5 | Pivot gene | oocyte/4-cell/8-cell/blastocyst |
| HSF2 | T5 | Pivot gene | 4-cell/8-cell/blastocyst |
| UBE3A | T5 | Pivot gene | oocyte/4-cell/blastocyst |
| RPS24 | T5 / CC3 | Pivot gene | oocyte/4-cell/8-cell/blastocyst |
| PRKCB | S2 | Intra-modular gene | 4-cell |
| RPL19 | S2 | Pivot gene | 4-cell/8-cell/blastocyst |
| SRPK1 | S2 | Pivot gene | 4-cell/8-cell/blastocyst |

We selected the T5 correlated gene UBE3A for a case study. In the co-expression network, UBE3A is a pivot gene crosstalking with three embryonic development stage specific modules for oocyte, 4-cell and blastocyst stage, respectively (Figure 6). The crosstalk between UBE3A and the oocyte/4-cell specific modules suggest potential underlying mechanisms explaining the key role of the T5 parameter in the early stage of preimplantation embryonic development, while the crosstalk with the blastocyst specific module helps us understand the impacts of the T5 parameter on blastocysts at the molecular level.

**Figure 6.**
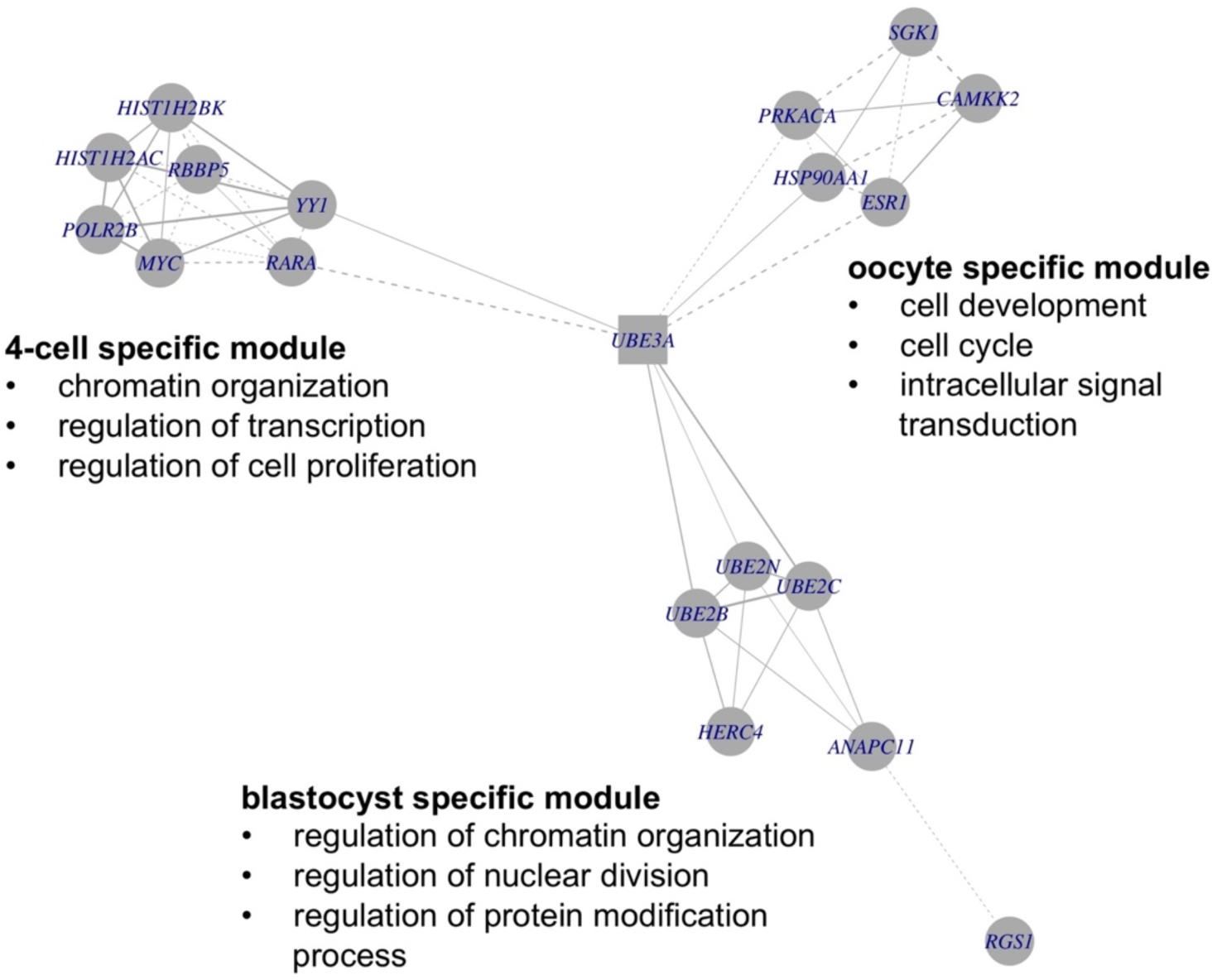
Crosstalk between pivot gene UBE3A and embryonic development stage specific modules. Nodes in square represent pivot genes and nodes in circle represent genes within modules. Edges in solid lines represent positive co-expression and edges in dashed lines represent negative co-expression.

## Discussion

This pre-clinical study showed that the transcriptome of human blastocysts which developed in a highly controlled, undisturbed culture environment with remote imaging via TL technology, is very similar to that of blastocysts grown in a standard box incubator system. It provided important reassurance that this IVF technology was not associated with fundamental changes in embryonic development, and allowed us to proceed with a clinical validation study prior to using the EmbryoScope routinely, in line with the model proposed by Kalleas et al (5). We then chose to examine the basis behind the use of TL imaging in predicting embryo developmental competence, by using gene co-expression analysis and system biology approaches to demonstrate that TL morphokinetic parameters associated with embryo developmental competence have correlates in the blastocyst transcriptome. This is important in furthering our understanding of the molecular biology of early embryonic development and for hypothesis-based approaches to developing TL embryo selection algorithms for clinical use.

### No fundamental changes of gene expression in undisturbed human embryo culture

In the statistical analysis of the ES and SI blastocysts gene expression data, we used very strict thresholds for filtering the high-confidence genes expressed in the two blastocyst groups to keep their fundamental gene expression patterns. In the unsupervised hierarchical clustering heatmap and PCA plot, we observed the ES and SI embryos were not classified into two distinct clusters based on the blastocysts gene expression, which suggests the ES blastocysts did not exhibit a different pattern in fundamental gene expression from the SI blastocysts. This analysis is consistent with the non-significant difference in quality between SI and ES blastocysts as Sciorio et al. reported (42). Sciorio et al. did report significantly higher proportions of good quality embryos in the ES group compared to the SI group on both day 2 and day 3, but not by blastocyst stage (42). In our subsequent clinical validation study (5), we also noted an increase in embryo cell number in ES compared to SI embryos on day 2 and 3, with ES embryos having similar if marginally lower grades for blastomere evenness and fragmentation. Given that there is some evidence that following transfer, ES embryos are more viable than SI (2–5), we conclude that any improvement in quality is negated by extended culture to blastocyst, or that quality differences relevant to improved embryo viability are not detected by our unsupervised clustering and PCA analysis. Nonetheless, the unsupervised clustering and PCA analysis provides reassurance that ES culture does not result in fundamental changes in embryonic biology.

In the differential gene expression analysis, it is to be noted that 40 (1.4%) out of the 2920 genes were identified as DEGs that were significantly upregulated or downregulated in ES compared to SI blastocysts. This represents a very small proportion, and clearly the differential gene expression did not change the fundamental gene expression patterns of ES blastocysts. However, the DEGs might provide clues for future studies to explore potential mechanisms in undisturbed human embryo culture.

### TL parameter correlated genes provide molecular links between the morphokinetic parameters and IVF clinical outcomes

Having established that the ES incubator system requirement to collect TL measurements did not in itself alter embryonic gene expression, we next explored the potential molecular pathways and genes underlying the association between TL parameters and embryonic development. We developed the network-constrained regression method eGBL, which provided superior performance in terms of feature gene selection in our simulation studies. We further focused on early markers of the degree of synchrony in embryonic progression (CC2, S2 and CC3, which respectively measure the duration of the 2-cell stage, 3-cell stage, and from 3-5 cell stage), along with T5 which marks the time taken to reach the 5-cell stage from fertilisation. We were able to identify TL parameter correlated genes, with some interesting biological insights which facilitate our understanding of the molecular links between the TL parameters and blastocyst development.

We found that TL parameter correlated genes were able to distinguish blastocysts with higher qualities and higher implantation potential (i.e. the good-quality group), since potential morphokinetic categories tended to be grouped together by the expression of TL parameter correlated genes. This finding helps to validate the clinical association between TL parameters and embryonic development competence, and further suggests a potential molecular level explanation which manifests in the blastocyst. We found 5 TL parameter correlated genes, CDX2, GRHL2, HAND1, PARP1 and PPARD, which are peri-implantation development marker genes and elicit important functions in trophectoderm development, trophoblast differentiation and placentation. The CC2 negative-correlated gene CDX2 is the core transcription factor responsible for trophectoderm development (39). The S2 negative-correlated gene HAND1 is essential for trophoblast differentiation and blastocyst implantation (37). The S2 negative-correlated gene GRHL2 is also a key regulator controlling trophoblast development (38), and the CC2 positive-correlated gene PARP1 has also been reported to be correlated with trophoblast differentiation (40). The CC3 positive-correlated gene PPARD plays important roles in placenta development (41). Trophoblast differentiation and placentation are the critical developmental processes required for the further development and nurturing of the blastocyst through the peri-implantation period. Dysfunction of these genes may result in developmentally compromised blastocysts, trophoblast and eventually placentae, which will reduce the success rate during IVF treatment and have implications for offspring health. These associations between early TL parameters and peri-implantation development marker genes might underlie the association between morphokinetic parameters and the success of embryonic implantation and clinical pregnancy. Thus, it is significant that three of these genes (CDX2, HAND1 and PPARD) were expressed at significantly different levels between the good-quality and poor-quality embryos.

### TL parameter correlated genes crosstalk with preimplantation embryonic development related gene functional modules

We next explored crosstalk between TL parameter correlated genes and the key development gene modules in our previously published gene co-expression network of human embryonic development (8). The pivot genes which link modules across several embryonic developmental stages (8) are of particular interest. While some are examples of housekeeping functions, e.g. the ribosomal proteins RPL19 and RPS24 and heat shock transcription factor 2 (HSF2), others may have important roles in the regulation of embryonic development.

We present the example of the T5 positive-correlated ubiquitin ligase UBE3A, which is involved in the process of targeting proteins for turnover via ubiquitin-tagged degradation. The identification of UBE3A as a pivot gene is consistent with our previous analysis of our human embryo gene co-expression networks, in which ubiquitin was identified as a key network module across a number of developmental stages (8). UBE3A specifically has been previously shown to be expressed throughout human preimplantation development (43, 44), and intriguingly, ubiquitin itself was detected as a major component of the human embryo secreted proteome (45). UBE3A has a number of specific target proteins including p53 (46), which has clearly established roles in regulating DNA repair and cell proliferation in preimplantation embryos (47) and UBE3A has been proven a critical gene to embryo implantation in mouse (48). Intriguingly, UB3EA is also paternally imprinted during neuronal development and loss of imprinting or mutation in the maternal UBE3A allele has been implicated in the related imprinted gene disorders Angelman and Prader-Willi syndromes, as well as autism, all of which are increased in ART offspring (49, 50). As a pivot gene, UBE3A crosstalks with multiple development stage specific gene modules including oocyte stage, 4-cell stage and blastocyst stage. Such crosstalk provide clues for potential molecular mechanisms associated with T5 which is a key parameter used for morphokinetic embryo selection as in the embryo classification methods proposed by Cruz et al. (11).

The pivot gene EXO1, also T5 positive-correlated, is an exonuclease with roles in DNA mismatch repair, telomere maintenance and homologous recombination during oocyte meiosis 1 (51). In embryonic development it mediates the response to environmental insults which cause DNA double strand breaks (52). Its role in regulating modules at all developmental stages from oocyte through to blastocyst clearly suggests the importance of these pathways at multiple points in early development.

Both UBE3A and EXO1 are T5 positive-correlated genes, as is EGFR, a developmentally vital gene associated with the 4-cell stage module. As T5 is the only marker we included which measures developmental progress rather than embryonic synchrony (CC2, S2 and CC3), this suggests that the developmentally important genes with regulatory roles throughout early development (of which EXO1 and UBE3A are examples) may be strongly associated with the developmental progress of the embryo. Conversely, these observations provide potential biological insights underpinning the predictive value of T5 for development of the embryo to clinical pregnancy.

It is worth noting that S2 was negatively correlated with two protein kinases, PRKCB and SRPK1, suggesting a role for cellular signaling processes in the regulation of the embryonic second cell cycle.

It is to be noted that pivot genes are enriched in T5 correlated genes and S2 correlated genes, but not CC2 and CC3. Although there is one CC3 negative-correlated gene, RPS24, identified as a pivot gene, it is a T5 negative-correlated gene as well. From biological perspectives, the developmental importance of T5 and S2 correlated genes suggest stronger associations of T5 and S2 with the blastocyst gene expression, which might suggest more crucial roles of T5 and S2 for morphokinetic embryo selection in clinical IVF treatments.

### Limitations of this study

This study was limited by the sample size of embryo transcriptomes. The unbalanced sample sizes of the time-lapse incubator developed embryos and the reference conventional incubator developed embryos might reduce the power and reproducibility in the statistical analysis of the ES and SI blastocysts gene expression. The limited sample size of the TL EmbryoScope incubator developed embryos might result in false positives and lower sensitivity in identification of TL parameter correlated genes because of overfitting in regression models. In the co-expression network analysis, pivot genes and gene modules were identified based on the gene-gene interaction network. The incompleteness of gene-gene interaction network might result in loss of pivot genes because the interactions between the genes and modules have not been identified yet. Larger sample size of embryo transcriptomes derived by advanced technologies (e.g., RNA-seq) and higher coverage of gene-gene interaction data will improve the sensitivity and robustness of this study.

## Conclusions

In this paper, we performed a systematic study of the gene expression and TL morphokinetic parameters in undisturbed human embryo culture. We validated that there is no fundamental difference in the developmental program of the embryos grown in the EmbryoScope time-lapse system, which provides important reassurance over the time-lapse technology in IVF. The TL parameter correlated genes, identified by network-constrained regression, suggest biological insights into the predictive capabilities of the TL parameters during preimplantation embryonic development (T5, S2, CC2 and CC3) for subsequent implantation and clinical pregnancy. The crosstalk between TL parameter correlated genes and the key gene functional modules in the human preimplantation embryonic development gene co-expression network provide clues for exploring further potential mechanisms during preimplantation development which might be associated with the TL parameter at the molecular level. This study suggests methodologies for assessing the impact of additional predictive correlates of value for optimal embryo development and survival in ART technologies, and understanding their mechanisms of action, especially in light of the added benefit of minimally disturbed culture conditions the TL EmbryoScope Incubator and its microwell chambers provide, limiting exposure of human IVF embryos to additional stress factors.

## Conflict of Interest

The authors declare that the research was conducted in the absence of any commercial or financial relationships that could be construed as a potential conflict of interest.

## Author Contributions

DB and PL supervised this research. DB, SK, GH and AD contributed to conceiving and designing the study. MK participated in the study design and contributed to data acquisition. PL, HX, AS, HS and KS contributed to data analysis and data interpretation. HX and DB wrote the manuscript. All authors contributed to the critical revisions of the manuscript and approved the final version of the manuscript.

## Acknowledgments

The work presented here has received funding from the European Union’s FP7 and Horizon 2020 Research and Innovation Programmes under the Marie Skłodowska-Curie Grant Agreement PITN-GA-2012-317146 (EPIHEALTHNET) and No. 812660 (DohART-NET) and grant agreements FP7-HEALTH-2011-TWO-STAGE-278418 (EPIHEALTH). In accordance with FP7 and H2020 rules, no new human embryos were sacrificed for research activities performed from the EU funding, as it concerned only in silico analyses of recorded time-lapse and transcriptomics datasets. We also acknowledge funding from the Medical Research Council (MR/L004992/1), Manchester NHS Foundation Trust, the NIHR Manchester Clinical Research Facility and the University of Manchester. The EmbryoScope used in the study was provided on loan by Unisense FertiliTech.

## Notes

### Competing Interest Statement

The authors have declared no competing interest.

## References

1. Mandal A. 8 million babies born through IVF says study. In: https://www.news-medical.net/news/20180704/8-million-babies-born-through-IVF-says-study.aspx, 2018.

2. Meseguer M, Herrero J, Tejera A, Hilligsoe KM, Ramsing NB, Remohi J. The use of morphokinetics as a predictor of embryo implantation. Human reproduction (Oxford, England) 2011;26:2658–71.

3. Meseguer M, Rubio I, Cruz M, Basile N, Marcos J, Requena A. Embryo incubation and selection in a time-lapse monitoring system improves pregnancy outcome compared with a standard incubator: a retrospective cohort study. Fertil Steril 2012;98:1481–9.e10.

4. Rubio I, Galan A, Larreategui Z, Ayerdi F, Bellver J, Herrero J et al. Clinical validation of embryo culture and selection by morphokinetic analysis: a randomized, controlled trial of the EmbryoScope. Fertil Steril 2014;102:1287–94.e5.

5. Kalleas D, McEvoy K, Horne G, Roberts SA, Brison DR. Live birth rate following undisturbed embryo culture at low oxygen in a time-lapse incubator compared to a high-quality benchtop incubator. Hum Fertil (Camb) 2020:1–7.

6. Armstrong S, Bhide P, Jordan V, Pacey A, Marjoribanks J, Farquhar C. Time-lapse systems for embryo incubation and assessment in assisted reproduction. The Cochrane database of systematic reviews 2019;5:Cd011320.

7. Harper J, Magli MC, Lundin K, Barratt CL, Brison D. When and how should new technology be introduced into the IVF laboratory? Human reproduction (Oxford, England) 2012;27:303–13.

8. Smith HL, Stevens A, Minogue B, Sneddon S, Shaw L, Wood L et al. Systems based analysis of human embryos and gene networks involved in cell lineage allocation. BMC genomics 2019;20:171.

9. Aguilar J, Motato Y, Escriba MJ, Ojeda M, Munoz E, Meseguer M. The human first cell cycle: impact on implantation. Reprod Biomed Online 2014;28:475–84.

10. Chamayou S, Patrizio P, Storaci G, Tomaselli V, Alecci C, Ragolia C et al. The use of morphokinetic parameters to select all embryos with full capacity to implant. J Assist Reprod Genet 2013;30:703–10.

11. Cruz M, Garrido N, Herrero J, Perez-Cano I, Munoz M, Meseguer M. Timing of cell division in human cleavage-stage embryos is linked with blastocyst formation and quality. Reprod Biomed Online 2012;25:371–81.

12. Alpha Scientists in Reproductive M, Embryology ESIGo. The Istanbul consensus workshop on embryo assessment: proceedings of an expert meeting. Human reproduction (Oxford, England) 2011;26:1270–83.

13. Brady G, Iscove NN. Construction of cDNA libraries from single cells. Methods in enzymology 1993;225:611–23.

14. Bloor DJ, Metcalfe AD, Rutherford A, Brison DR, Kimber SJ. Expression of cell adhesion molecules during human preimplantation embryo development. Molecular human reproduction 2002;8:237–45.

15. Shaw L, Sneddon SF, Zeef L, Kimber SJ, Brison DR. Global gene expression profiling of individual human oocytes and embryos demonstrates heterogeneity in early development. PloS one 2013;8:e64192.

16. RCoreTeam. R: A language and environment for statistical computing. R Foundation for Statistical Computing, Vienna, Austria https://wwwr-projectorg/2016.

17. Gentleman RC, Carey VJ, Bates DM, Bolstad B, Dettling M, Dudoit S et al. Bioconductor: open software development for computational biology and bioinformatics. Genome biology 2004;5:R80.

18. Kauffmann A, Gentleman R, Huber W. arrayQualityMetrics--a bioconductor package for quality assessment of microarray data. Bioinformatics (Oxford, England) 2009;25:415–6.

19. Irizarry RA, Hobbs B, Collin F, Beazer-Barclay YD, Antonellis KJ, Scherf U et al. Exploration, normalization, and summaries of high density oligonucleotide array probe level data. Biostatistics (Oxford, England) 2003;4:249–64.

20. Pepper SD, Saunders EK, Edwards LE, Wilson CL, Miller CJ. The utility of MAS5 expression summary and detection call algorithms. BMC bioinformatics 2007;8:273.

21. Gautier L, Cope L, Bolstad BM, Irizarry RA. affy--analysis of Affymetrix GeneChip data at the probe level. Bioinformatics (Oxford, England) 2004;20:307–15.

22. Ritchie ME, Phipson B, Wu D, Hu Y, Law CW, Shi W et al. limma powers differential expression analyses for RNA-sequencing and microarray studies. Nucleic acids research 2015;43:e47.

23. Benjamini Y, Drai D, Elmer G, Kafkafi N, Golani I. Controlling the false discovery rate in behavior genetics research. Behavioural brain research 2001;125:279–84.

24. TheGeneOntologyConsortium. Expansion of the Gene Ontology knowledgebase and resources. Nucleic acids research 2017;45:D331–d8.

25. Cerami EG, Gross BE, Demir E, Rodchenkov I, Babur O, Anwar N et al. Pathway Commons, a web resource for biological pathway data. Nucleic acids research 2011;39:D685–90.

26. Pan W, Xie B, Shen X. Incorporating predictor network in penalized regression with application to microarray data. Biometrics 2010;66:474–84.

27. Becker E, Robisson B, Chapple CE, Guenoche A, Brun C. Multifunctional proteins revealed by overlapping clustering in protein interaction network. Bioinformatics (Oxford, England) 2012;28:84–90.

28. Subramanian A, Tamayo P, Mootha VK, Mukherjee S, Ebert BL, Gillette MA et al. Gene set enrichment analysis: a knowledge-based approach for interpreting genome-wide expression profiles. Proceedings of the National Academy of Sciences of the United States of America 2005;102:15545–50.

29. Breheny P. The group exponential lasso for bi-level variable selection. Biometrics 2015;71:731–40.

30. Tibshirani R. Regression shrinkage and selection via the lasso. Journal of the Royal Statistical Society Series B (Methodological) 1996:267–88.

31. Zou H, Hastie T. Regularization and Variable Selection via the Elastic Net. Journal of the Royal Statistical Society: Series B (Statistical Methodology) 2005;67:301–20.

32. Li C, Li H. Network-constrained regularization and variable selection for analysis of genomic data. Bioinformatics (Oxford, England) 2008;24:1175–82.

33. Gil-Cayuela C, Lopez A, Martinez-Dolz L, Gonzalez-Juanatey JR, Lago F, Rosello-Lleti E et al. The altered expression of autophagy-related genes participates in heart failure: NRBP2 and CALCOCO2 are associated with left ventricular dysfunction parameters in human dilated cardiomyopathy. PloS one 2019;14:e0215818.

34. Wilkerson DC, Murphy LA, Sarge KD. Interaction of HSF1 and HSF2 with the Hspa1b promoter in mouse epididymal spermatozoa. Biol Reprod 2008;79:283–8.

35. Bai S, Grossman G, Yuan L, Lessey BA, French FS, Young SL et al. Hormone control and expression of androgen receptor coregulator MAGE-11 in human endometrium during the window of receptivity to embryo implantation. Molecular human reproduction 2008;14:107–16.

36. Richani D, Gilchrist RB. The epidermal growth factor network: role in oocyte growth, maturation and developmental competence. Hum Reprod Update 2018;24:1–14.

37. Scott IC, Anson-Cartwright L, Riley P, Reda D, Cross JC. The HAND1 basic helix-loop-helix transcription factor regulates trophoblast differentiation via multiple mechanisms. Mol Cell Biol 2000;20:530–41.

38. Walentin K, Hinze C, Werth M, Haase N, Varma S, Morell R et al. A Grhl2-dependent gene network controls trophoblast branching morphogenesis. Development 2015;142:1125–36.

39. Strumpf D, Mao CA, Yamanaka Y, Ralston A, Chawengsaksophak K, Beck F et al. Cdx2 is required for correct cell fate specification and differentiation of trophectoderm in the mouse blastocyst. Development 2005;132:2093–102.

40. Hemberger M, Nozaki T, Winterhager E, Yamamoto H, Nakagama H, Kamada N et al. Parp1-deficiency induces differentiation of ES cells into trophoblast derivatives. Dev Biol 2003;257:371–81.

41. Barak Y, Sadovsky Y, Shalom-Barak T. PPAR Signaling in Placental Development and Function. Ppar Res 2008;2008:142082.

42. Sciorio R, Thong JK, Pickering SJ. Comparison of the development of human embryos cultured in either an EmbryoScope or benchtop incubator. J Assist Reprod Genet 2018;35:515–22.

43. Salpekar A, Huntriss J, Bolton V, Monk M. The use of amplified cDNA to investigate the expression of seven imprinted genes in human oocytes and preimplantation embryos. Molecular human reproduction 2001;7:839–44.

44. Monk M, Salpekar A. Expression of imprinted genes in human preimplantation development. Mol Cell Endocrinol 2001;183 Suppl 1:S35–40.

45. Katz-Jaffe MG, Schoolcraft WB, Gardner DK. Analysis of protein expression (secretome) by human and mouse preimplantation embryos. Fertil Steril 2006;86:678–85.

46. Scheffner M, Huibregtse JM, Vierstra RD, Howley PM. The HPV-16 E6 and E6-AP complex functions as a ubiquitin-protein ligase in the ubiquitination of p53. Cell 1993;75:495–505.

47. Wilson Y, Morris ID, Kimber SJ, Brison DR. The role of Trp53 in the mouse embryonic response to DNA damage. Molecular human reproduction 2019;25:397–407.

48. Wang H, Dey SK. Roadmap to embryo implantation: clues from mouse models. Nat Rev Genet 2006;7:185–99.

49. Gosden R, Trasler J, Lucifero D, Faddy M. Rare congenital disorders, imprinted genes, and assisted reproductive technology. Lancet (London, England) 2003;361:1975–7.

50. Liu L, Gao J, He X, Cai Y, Wang L, Fan X. Association between assisted reproductive technology and the risk of autism spectrum disorders in the offspring: a meta-analysis. Scientific reports 2017;7:46207.

51. Qiu J, Qian Y, Chen V, Guan MX, Shen B. Human exonuclease 1 functionally complements its yeast homologues in DNA recombination, RNA primer removal, and mutation avoidance. The Journal of biological chemistry 1999;274:17893–900.

52. Rein K, Yanez DA, Terre B, Palenzuela L, Aivio S, Wei K et al. EXO1 is critical for embryogenesis and the DNA damage response in mice with a hypomorphic Nbs1 allele. Nucleic acids research 2015;43:7371–87.

