## Supplementary Figures for "Interaction between gene expression and morphokinetic parameters in undisturbed human embryo culture"

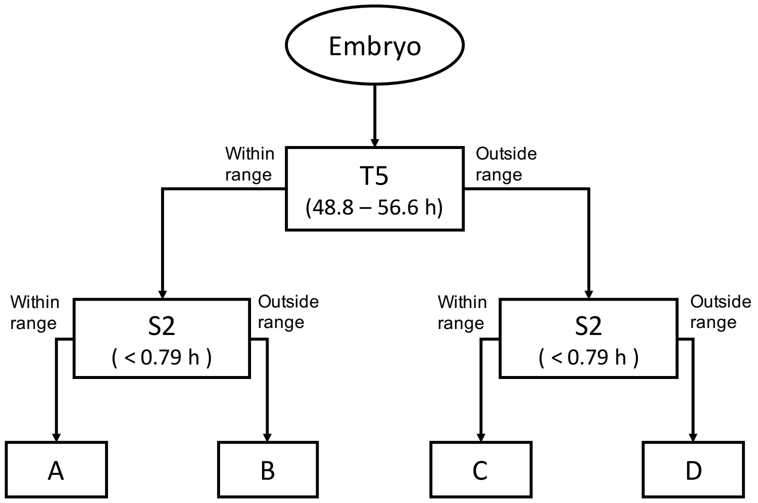


**Figure S1** Morphokinetic categories of embryos based on T5 and S2 proposed by Cruz et al. The numbers shown in parentheses are the optimal ranges for the corresponding TL parameters, i.e., 48.8-56.6 hours for T5 and less than 0.79 hours for S2 as the optimal ranges. Embryos were first bi-classified as within or outside the T5 optimal range. For the group within T5 range, embryos within or outside the S2 optimal range were classified as category A or B, respectively. Similarly, embryos in the group outside the T5 range were classified as category C or D.

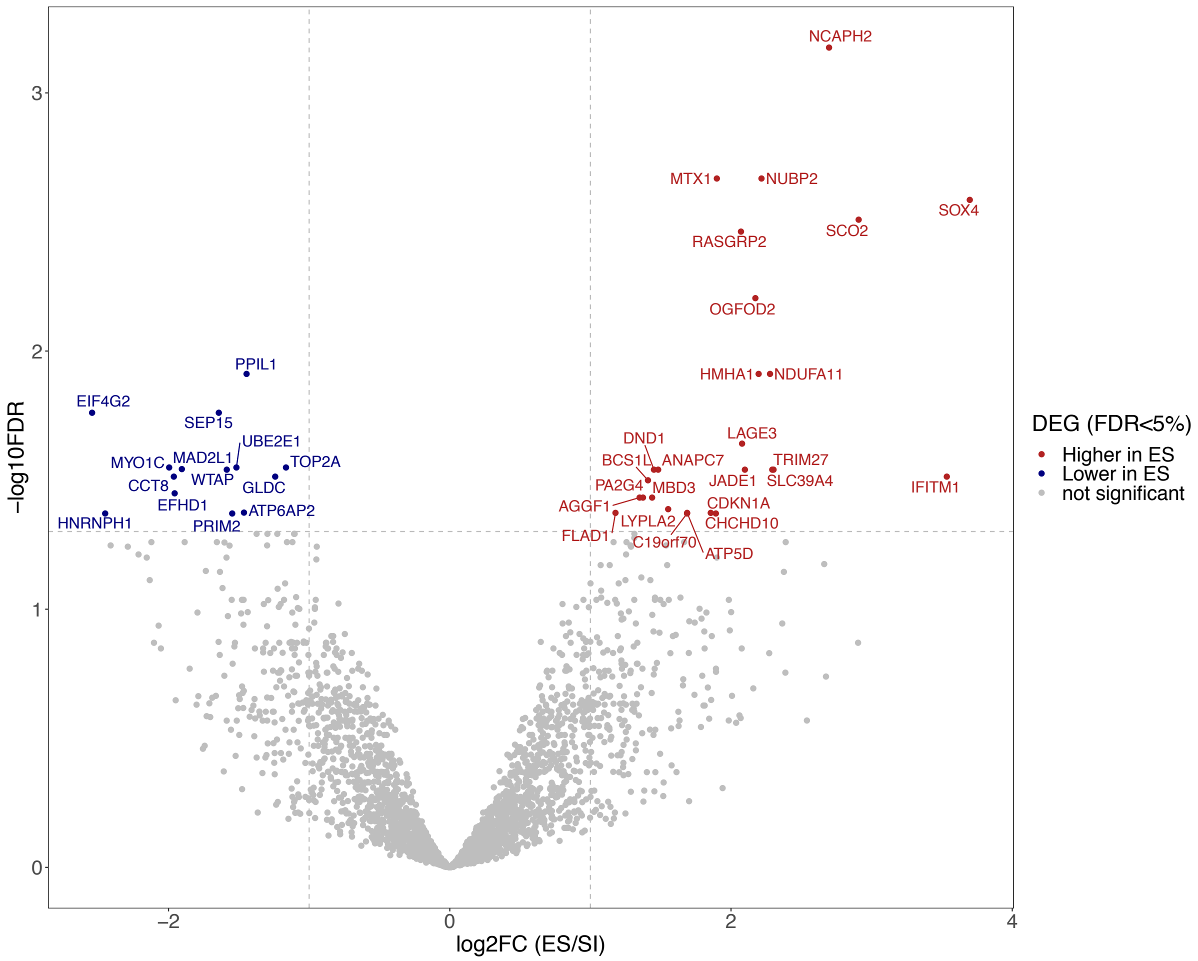


**Figure S2** Gene expression fold changes and significances in ES blastocysts compared to SI blastocysts.

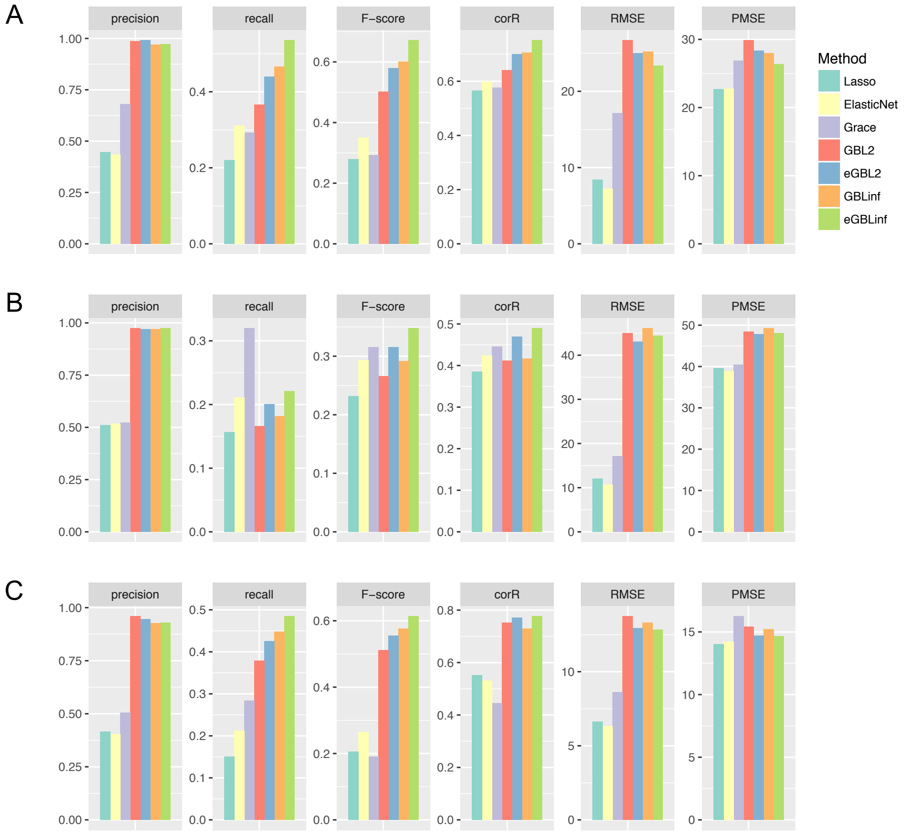


**Figure S3** Performances of regression methods on simulation studies. (A) Simulation study 1: A weighted gene co-expression network compromised of 50 subnetworks, each of which includes one gene hub interacting with 10 neighbouring genes with the corresponding decreasing co-expression weights as w = (0.95,0.85,...,0.05). The expression levels for the 50 hub genes in the simulated network exhibit standard normal distributions and the expression levels of its neighbour genes exhibit normal distributions with correlations w to standard normal. 10% of the genes were assumed as features that are related to the response variable and their neighbour genes were simulated with non-zero coefficients. The training set and test set were simulated in the same way with 50 samples and 100 samples respectively. (B) Simulation study 2: Simulated data were set up in the same way as simulation study 1, but with the assumption that 20% of the genes are features. (C) Simulation study 3: Simulated data were set up in the same way as simulation study 1, but assuming 30% of the coefficients with opposite signs for the neighbour genes of each gene hub. The regression methods include the proposed network constrained regression method eGBL with 2 and $\infty$ for parameter $\gamma$, two network constrained regression methods Grace and GBL, and two common regularised regression methods Lasso and ElasticNet. Evaluation measures include: The Precision measure referring to the proportion of the predicted positive samples are correct; The Recall measure referring to the proportion of the actual positive samples which are identified correctly; The F-score measure which is the harmonic mean of precision and recall; The Correlation (corR) between the estimated coefficients and the actual coefficients; The Root Mean Squared Error (RMSE), which is the square root of the mean of the squared errors on the training set; The Prediction Mean Squared Error (PMSE), which is the square root of the mean of the squared errors on the independent test set.

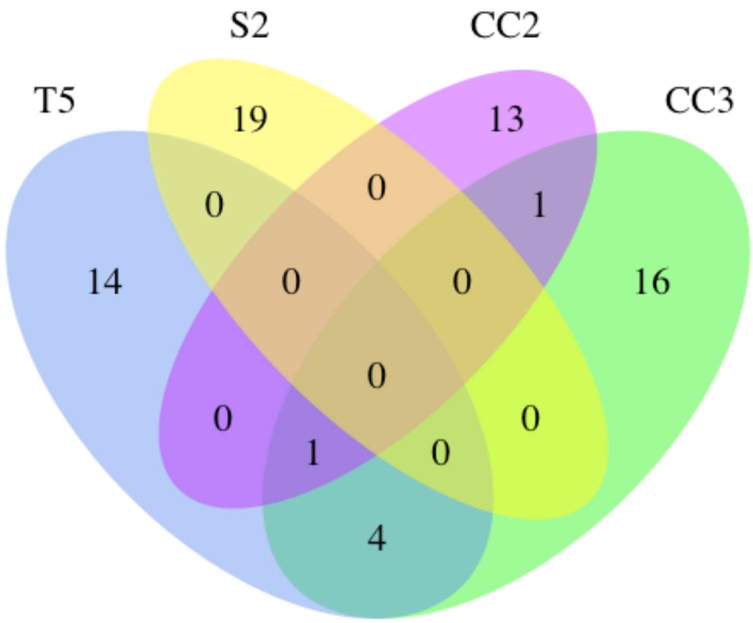


**Figure S4** Overlap between TL parameter correlated genes

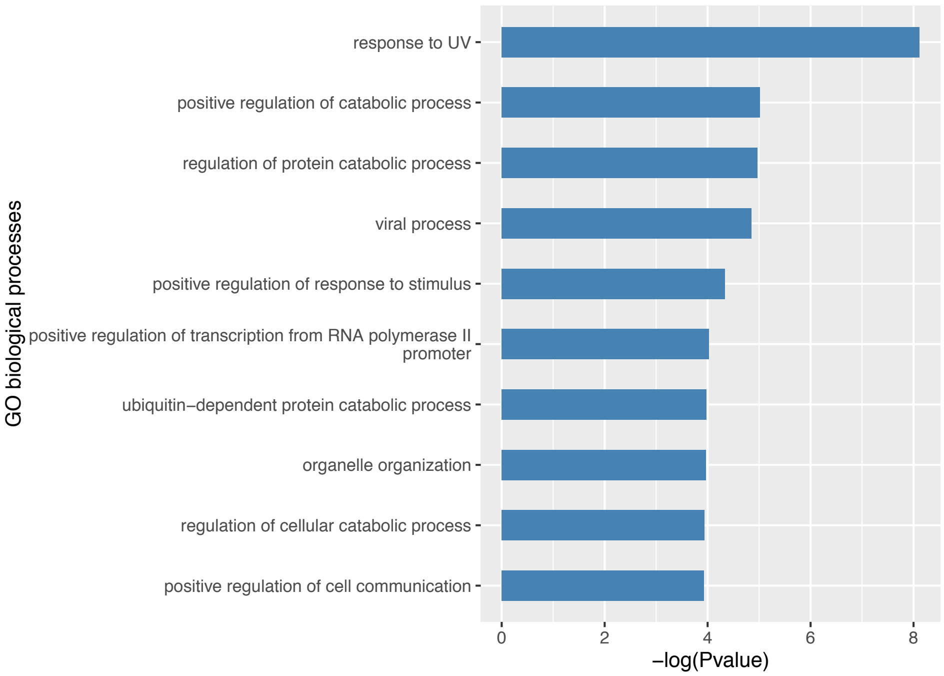


**Figure S5** Gene Ontology biological processes enriched by the time-lapse parameter T5 correlated genes.

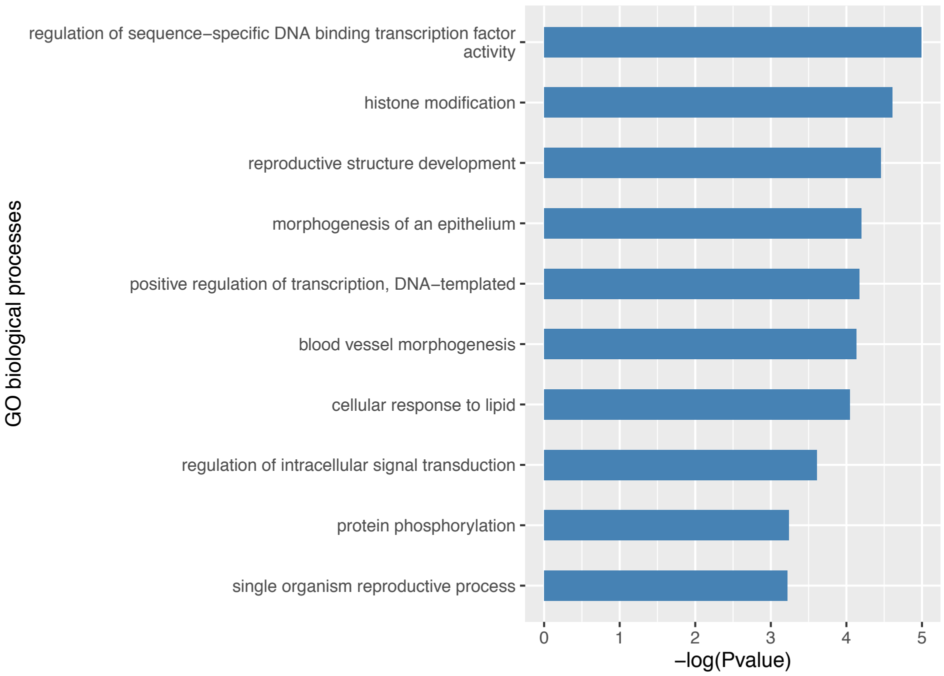


**Figure S6** Gene Ontology biological processes enriched by the time-lapse parameter S2 correlated genes.

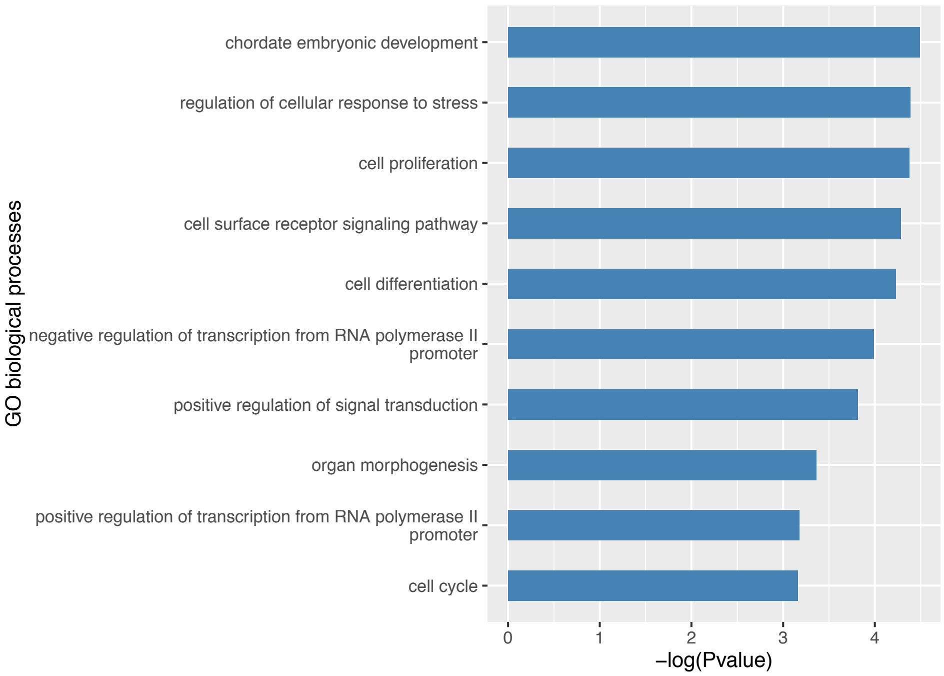


**Figure S7** Gene Ontology biological processes enriched by the time-lapse parameter CC2 correlated genes.

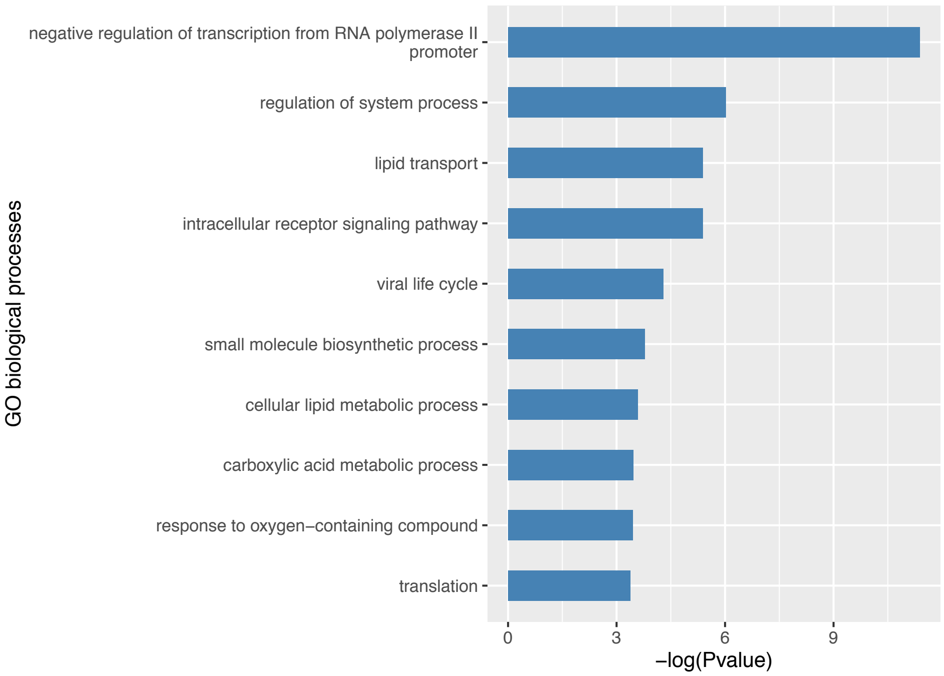


**Figure S8** Gene Ontology biological processes enriched by the time-lapse parameter CC3 correlated genes.
